# Towards a general model for predicting minimal metal concentrations co-selecting for antibiotic resistance plasmids

**DOI:** 10.1101/2020.09.14.295766

**Authors:** Sankalp Arya, Alexander Williams, Saul Vazquez Reina, Charles W. Knapp, Jan-Ulrich Kreft, Jon L. Hobman, Dov J. Stekel

## Abstract

Many antibiotic resistance genes co-occur with resistance genes for transition metals, such as copper, zinc, or mercury. In some environments, a positive correlation between high metal concentration and high abundance of antibiotic resistance genes has been observed, suggesting co-selection due to metal presence. Of particular concern is the use of copper and zinc in animal husbandry, leading to potential co-selection for antibiotic resistance in animal gut microbiomes, slurry, manure, or amended soils. For antibiotics, predicted no effect concentrations have been derived from laboratory measured minimum inhibitory concentrations and some minimal selective concentrations have been investigated in environmental settings. However, minimal co-selection concentrations for metals are difficult to identify. Here, we use mathematical modelling to provide a general mechanistic framework to predict minimal co-selective concentrations for metals, given knowledge of their toxicity at different concentrations. We apply the method to copper (Cu), Zinc (Zn), mercury (Hg), lead (Pb) and silver (Ag), predicting their minimum co-selective concentrations in mg/L (Cu: 5.5, Zn: 1.6, Hg: 0.0156, Pb: 21.5, Ag: 0.152). To exemplify use of these thresholds, we consider metal concentrations from slurry and slurry-amended soil from a UK dairy farm that uses copper and zinc as additives for feed and antimicrobial footbath: the slurry is predicted to be co-selective, but not the slurry-amended soil. This modelling framework could be used as the basis for defining standards to mitigate risks of antimicrobial resistance applicable to a wide range of environments, including manure, slurry and other waste streams.

## Introduction

The persistence and spread of antimicrobial resistance (AMR) is a major global threat, with at least 700,000 deaths per year attributed to bacterial infections by drug-resistant strains world-wide [1]. Reduction of antibiotic use, or cessation of use of some veterinary antibiotics, is seen as critically important to mitigate the threat of AMR. A classic example of this strategy has been the banning of avoparcin in poultry production, e.g. Germany and Denmark in 1995, other EU countries by 1997 and Taiwan in 2000. The success of this ban can be exemplified with Norwegian poultry farms showing high abundance (99%) of vancomycin-resistant enterococci (VRE) in farms exposed to avoparcin prior to the ban and lower abundance (11%) in samples from unexposed farms [2], while in Taiwan there was a decrease from 25% farms having vancomycin-resistant enterococci (VRE) in 2000 to 8.8% farms in 2003 [3].

However, the continued presence of resistant strains suggests that there may be other factors that promote persistence of antibiotic resistance genes (ARGs). One of these factors is co-selection: selective pressure exerted by a toxicant that maintains other ARGs. This can occur in different ways: (i) co-resistance, i.e., multiple genes encoding resistance against different antibiotics and metals which are on the same genetic element; (ii) cross-resistance, i.e., the same mechanism (e.g., efflux pumps) providing resistance against multiple toxicants; (iii) co-regulation, which is the coordinated response to the presence of either antibiotic or metal, this activates mechanisms necessary for the resistance against the other or both [4]. High metal concentration provides co-selective pressure for certain antibiotics. For example, Lee *et al.* (2005) showed that the *mdt* operon, which encodes for a multidrug resistance efflux pump in *E. coli* was up-regulated in response to excess zinc [5]. Resistance to antibiotics is also enriched in response to metal shock loading [6, 7], or due to long-term exposure to metal [8, 9]. However, most data only provides indirect evidence co-selection due to metal presence, by providing evidence of co-occurrence of metal and antibiotic resistance, in many environments, including oral and intestinal [10], sludge bioreactors [11], marine [12] and soil sediments [13].

Selective pressure can be caused at concentrations lower than the minimal inhibitory concentration (MIC). The FAO and WHO support the concept of minimum selective concentration (MSC) for antibiotics, i.e., a threshold concentration above which the resistance genes are selected. MSCs are available for antibiotics, based on standard MICs, through both empirical and modelling approaches [14–17], although environmental studies show the issue to be much more complex [18]. However, co-selection pressure due to transition metals might mean that in some environments, ARGs could be selected for and maintained in a bacterial community even with antibiotic concentrations below MSC. Metal and antibiotic resistance genes co-occur in environments where the metal contamination [19, 20] is sufficiently high to provide co-selection pressure for persistence and proliferation of antibiotic resistance [10, 21–23]. The notion of Minimal Co-Selective Concentrations (MCSCs) for transition metals was introduced by Seiler and Berendonk [13], who identified possible thresholds based upon observations of metal concentrations in a range of environments. However, the lack of appropriate MCSCs has been highlighted by the FAO and WHO [24]; indeed a rigorous and consistent approach to defining MCSCs could be used, alongside toxicity, to inform suitable standards for metal concentrations in agriculture or environmental contexts.

We address this research gap using a mathematical modelling approach. Models can help to understand and predict the impact of co-selection under different scenarios, and have already helped in understanding factors associated with AMR emergence and spread such as mutation rates [25], antibiotic consumption [26], water troughs on farms [27] as well as quantifying the importance of factors such as conjugation [28]. Models have also accurately predicted MIC values of *β*-lactams against MRSA [29]. One of the few mathematical models for co-selection studied the concentration of resistant bacteria in the Poudre River in Colorado and determined that external input and selection pressure solely due to tetracycline was insufficient to explain the observed levels of resistant bacteria. A co-selection model, on the other hand, which considered both tetracycline and metal concentrations reproduced the observed data [30].

In this study, we developed a general model to quantify the effect of metal concentrations on persistence of resistance in a bacterial population, using an approach that can be applied to any metal and any environment, given knowledge of its toxicity in that environment. We compare the results from deterministic (applicable to large well-mixed populations) and stochastic versions (applicable to small populations where random events may be significant) of the same model. We then analyse the effects of metal toxicity and plasmid fitness cost on the persistence of resistance in each version of the model. The model allows us to identify MCSCs for transition metals and how they depend on the toxicity of the metal and the fitness cost of carrying resistance. We show that both deterministic and stochastic versions of the model provide similar results, with resistance lost only at high fitness costs and sufficiently dilute metal concentrations, i.e. low toxicity. However, the stochastic model does suggest a higher chance of persistence for several months without antibiotic selection. Finally, we demonstrate the use of the MSCSs by applying them to measurements of copper and zinc concentrations in dairy slurry and slurry-amended soil on the same farm.

## Materials and methods

### Model description

The purpose of this analysis is to understand how the persistence of resistance genes is dependent on the fitness cost of carrying the resistance genes (for example on a plasmid) and the selective pressure from metals being present in the environment. We also investigate how deterministic and stochastic modelling paradigms impact upon the results. The models describe a generalised process of conjugation transferring the resistance genes, how bacterial growth is affected by the fitness cost of plasmid carriage (if present), and how death is affected by the concentration of metal in the environment.

We model a small (micro-scale) volume element representative of a larger system. The complete macro-scale ecosystem can be considered to be made up of replicates of the modelled domain [31, 32]. The starting bacterial population is of primarily resistant bacteria (99.32%), without antibiotic present, because we are interested in the persistence of resistance rather than the spread of resistance. The deterministic version of the model is described by ordinary differential equations (ODEs) Eq (1)–(2). As in Baker *et al.* [28], bacterial growth is defined by a logistic growth terms, affected by the fitness cost (*α*) of the plasmid for the resistant cells. Conjugation uses a classic Sensitive-Infected-Resistant/Recovered (SIR) model formulation for plasmid transfer, with “infection” of sensitive cells (S) by resistant cells (R) with rate constant *β*. The differences from the Baker *et al.* model [28] are the inclusion of death of sensitive (*δ*_*S*_) and resistant bacteria (*δ*_*R*_) - at different rates due to the resistance to metal - and plasmid loss (*ϵ*) due to segregation upon cell division.

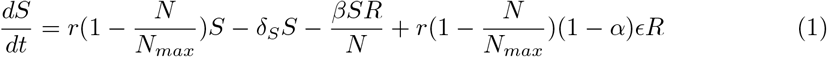

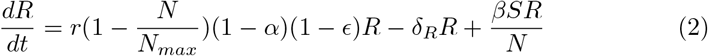

where *N* = *S* + *R*. The same model structure can be described by a set of discrete events which define the stochastic simulation algorithm (SSA). Table 1 provides the details of this SSA. Each event has a reaction rate which is the same as the rates defined in ODEs Eq (1)–(2).

**Table 1.** Stochastic Simulation Algorithm for Eq (1) – (2)

| Event | Description | Rate |
| --- | --- | --- |
| $S \rightarrow 2S$ | Growth of sensitive bacteria | $r(1 - \frac{N}{N_{max}})S$ |
| $S \rightarrow$ | Death of sensitive bacteria | $\delta_S S$ |
| $R \rightarrow 2R$ | Growth of resistant bacteria | $r(1 - \frac{N}{N_{max}})(1 - \alpha)R$ |
| $R \rightarrow$ | Death of resistant bacteria | $\delta_R R$ |
| $S \rightarrow R$ | Conjugation | $\frac{\beta SR}{N}$ |
| $R \rightarrow S + R$ | Plasmid loss due to segregation | $r(1 - \frac{N}{N_{max}})(1 - \alpha)\epsilon R$ |

The table lists the events describing the lifecycle of the bacteria as well as the processes of conjugation and plasmid loss. The rates for the different events are the same as in the ODE version of the model.

We estimated the parameters for bacterial death rate for both resistant and sensitive bacteria under different metal concentrations using the metal toxicity values for *E. coli* provided by Ivask *et al.* [33] and Equation (3). We used the SCO (Social Cognitive Optimization) evolutionary solver in LibreOffice Calc to estimate the unknown parameters *E*_*max*_ (maximum death rate due to metal), MIC (Minimum Inhibitory Concentration) and H (Hill coefficient) for all metals; fits to metal toxicity data are shown in Figure S5, demonstrating successful model calibration.

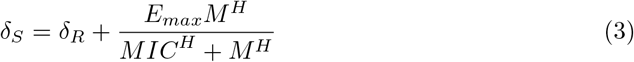

where, *M* is the metal concentration. Once the parameters were estimated, the concentrations of copper and zinc measured using the ICP-MS techniques, for both slurry and slurry treated soil, were used to calculate the ratio of death rate of sensitive to resistant bacteria (*ζ*).

### Numerical solutions of model

For the deterministic model, differential equations were simulated using the R [34] deSolve package [35] LSODA algorithm and sensitivity analysis was performed using the rootSolve [36, 37], doParallel [38] and foreach [39] R packages. For the parameter sensitivity analysis using the stochastic model, we used COPASI [40], and created shell scripts to run each parameter combination one thousand times. The output of each run was then imported into R to produce parameter sensitivity graphs with the ggplot2 [41] package, with 5-dimensional data expressed as two spatial dimensions and three colour dimensions, using an RGB combination for each point associated with each parameter combination, red for persistence of resistance, blue for loss of resistance, and green for total cell death. For example, if out of 1000 runs of the stochastic version of the model, 300 runs predicted persistence of resistance and 700 runs loss of resistance, then a colour 30% red and 70% blue would be plotted.

For example environments, we took the measured values of copper and zinc concentrations in slurry and slurry amended soil and calculated the death rate ratio of sensitive to resistant populations based on the metal toxicity values of *E. coli* [33], to check resistance fixation conditions in each environment, as described in the Model Overview section.

### Multi-elemental analysis by ICP-MS

#### Slurry

Slurry samples (2 mL) were acid digested on a hot plate using 6 mL Primar Plus grade HNO_3_ (68%) and 2 mL H_2_O_2_ (Thermo Fisher Scientific, Loughborough, UK). Samples were diluted with Milli-Q water (18.2 MΩ cm) to 50 mL and syringe filtered to <0.2 *μ*m (Merck-Millipore, Burlington, USA) prior to analysis by inductively coupled plasma mass spectrometry (icapQ model; Thermo Fisher Scientific, Bremen, Germany). Samples were introduced (flow rate 1.2 mL min^−1^) from an autosampler (Cetac ASX-520) incorporating an ASXpress rapid uptake module through a perfluoroalkoxy (PFA) Microflow PFA-ST nebuliser (Thermo Fisher Scientific, Bremen, Germany). Sample processing was undertaken using Qtegra software (Thermo-Fisher Scientific) utilizing external cross-calibration between pulse-counting and analogue detector modes when required. The ICP-MS was run employing two operational modes with in-sample switching between a collision cell (i) charged with He gas with kinetic energy discrimination (KED) to remove polyatomic interferences and (ii) using H_2_ gas as the cell gas. The latter was used only for Se determination. Peak dwell times were 100 ms with 150 scans per sample.

Internal standards, used to correct for instrumental drift, were introduced to the sample stream on a separate line (equal flow rate) via the ASXpress unit and included Sc (10*μ*g/L), Ge (10 *μ*g/L), Rh (5 *μ*g/L) and Ir (5 *μ*g/L). The matrix used for internal standards, calibration standards and sample dilution was 2% Primar Plus grade HNO_3_ with 4% methanol (to enhance ionization of some elements such as Se).

Calibration standards included (i) a multi-element solution with Ag, Al, As, Ba, Be, Cd, Ca, Co, Cr, Cs, Cu, Fe, K, Li, Mg, Mn, Mo, Na, Ni, P, Pb, Rb, S, Se, Sr, Ti, Tl, U, V and Zn, in the range 0 to 100 *μ*g/L (0, 20, 40, 100 *μ*g/L) (Claritas-PPT grade CLMS-2 from SPEX Certiprep Inc., Metuchen, NJ, USA); (ii) a bespoke external multi-element calibration solution (PlasmaCAL, SCP Science, France) with Ca, Mg, Na and K in the range 0-30 mg/L and (iii) a mixed phosphorus, boron and sulphur standard made in-house from salt solutions (KH_2_PO_4_, K_2_SO_4_ and H_3_BO_3_).

#### Soil

For extractable macro- and micro-elemental analysis, 1 g of soil was suspended in 9 mL of 1 M NH_4_NO_3_ and mixed thoroughly by agitation using a rotary shaker for 1 hour. Subsequently samples were centrifuged and 1 mL of the resulting supernatant was diluted in 9 mL of 2% nitric acid. Finally, samples were passed through a 0.22 *μ*m filter before being loaded for inductively coupled plasma mass spectrometry (ICP-MS; Thermo-Fisher Scientific iCAP-Q; Thermo Fisher Scientific, Bremen, Germany)

### Parameters used in the models

The parameters were taken to match the model parameters of Baker *et al.* [28], where possible; other parameter values were taken from references in Table 2. The estimated death rates for metals from the metal toxicity model are also listed. For sensitivity analysis, the ratio of death rates (*ζ*) and the fitness cost of carrying the plasmid with resistance genes are varied over a range. We also increased and decreased transfer frequency and probability of segregational loss, to see the effects of these two parameters on the output.

**Table 2.** All parameters used in Equations (1) – (3).

| Parameter | Description | Value (Range) | Source |
| --- | --- | --- | --- |
| $r$ | Specific growth rate | **0.5** h<sup>-1</sup> | [42–44] |
| $N_{max}$ | Carrying capacity of liquid slurry | $6.71 \times 10^7$ CFU/L | [45] |
| $\delta_R$ | Death rate of resistant bacteria (base rate) | **0.025** h<sup>-1</sup> | [46] |
| $\delta_S$ | Death rate of sensitive bacteria | $\zeta \cdot \delta_R$ | varied |
| $\zeta$ | Ratio of $\delta_S$ to $\delta_R$ affected by metal concentration | **1.0** (1.0-10.0) | varied |
| $\alpha$ | Fitness cost for carrying resistance as a fraction of $r$ | **0.1** (0-0.99) | [43, 47, 48] |
| $\beta$ | Conjugation rate | **0.001** h<sup>-1</sup> | [48, 49] |
| $\epsilon$ | Plasmid loss probability | **0.000144** | [50] |
| Volume | Volume of microcosm | $2.5 \times 10^{-6}$ L | Assumed |
| Copper (CuSO<sub>4</sub>) |  |  |  |
| MIC | Minimum inhibitory concentration | **212.79** mg/L | Estimated |
| $E_{max}$ | Maximum death rate due to metal | **1.74** h<sup>-1</sup> | Estimated |
| H | Hill coefficient | **1.54** | Estimated |
| Zinc (ZnSO<sub>4</sub>) |  |  |  |
| MIC | Minimum inhibitory concentration | **2760.31** mg/L | Estimated |
| $E_{max}$ | Maximum death rate due to metal | **1.37** h<sup>-1</sup> | Estimated |
| H | Hill coefficient | **0.72** | Estimated |
| Mercury (HgCl<sub>2</sub>) |  |  |  |
| MIC | Minimum inhibitory concentration | **1.85** mg/L | Estimated |
| $E_{max}$ | Maximum death rate due to metal | **5.89** h<sup>-1</sup> | Estimated |
| H | Hill coefficient | **1.44** | Estimated |
| Lead (Pb(NO<sub>3</sub>)<sub>2</sub>) |  |  |  |
| MIC | Minimum inhibitory concentration | **1728.7** mg/L | Estimated |
| $E_{max}$ | Maximum death rate due to metal | **18.74** h<sup>-1</sup> | Estimated |
| H | Hill coefficient | **1.82** | Estimated |
| Silver (AgNO<sub>3</sub>) |  |  |  |
| MIC | Minimum inhibitory concentration | **0.48** mg/L | Estimated |
| $E_{max}$ | Maximum death rate due to metal | **2.42** h<sup>-1</sup> | Estimated |
| H | Hill coefficient | **5.19** | Estimated |

## Results

In these simulations, we consider the persistence of the resistant strains following withdrawal of antibiotic, but in the continued presence of metal. Therefore the population starts at 99.3% resistant cells with only a small concentration of sensitive cells. For the stochastic models, we consider a microcosm of this population. We vary the ratio of death rates (sensitive/resistant) between 1.0, corresponding to an absence of toxicity, so sensitive and resistant cells die at the same rate, and thus there is no selection pressure; and 10.0, corresponding to strong selective pressure, with sensitive cells dying ten times faster than resistant cells due to metal toxicity. We vary the fitness cost between 0 (no fitness cost) and 1 (hosts carrying plasmid cannot grow). This full range of fitness costs is included for analytical completeness so that model behaviour can be fully understood; the typical biologically realistic range is from 0.1 to 0.3 [43, 47, 48].

### To persist or not to persist and the role of chance

To demonstrate model behaviour, we show model simulations for the four bounding values of fitness cost and death rate ratio used in the sensitivity analysis below. Thus all model behaviours in the sensitivity analysis lie in between these extremes. When the fitness cost is 0, the resistant bacteria persist in the population, irrespective of the death rate ratio (Fig 1(a) and (c)). When stochasticity is introduced into the model, then in the absence of selection (i.e. death rate ratio of 1) then a small proportion of simulations saw a loss of plasmid due to drift (2.6% of cases). Under strong selective conditions (death rate ratio of 10), the plasmid is fully maintained in both the deterministic and stochastic simulations. For extreme values of fitness cost (1), the proportion of resistance cells decreases over time. The rate of decrease depends upon the level of selective pressure. In the absence of selective pressure (death rate ratio of 1), resistance persists for approximately 40 days, before decreasing sharply (Fig 1(b)). Under strong selection pressure (death rate ratio of 10), resistance persists for longer, declining after about 80 days (Fig 1(d)). In the stochastic model under these conditions, both sensitive and resistant cells die, and no meaningful results can be shown. From these graphs, it can be inferred that under intermediate values of fitness cost and death rate ratio, resistance will persist for different periods of time. This is explored fully next.

**Fig 1.**
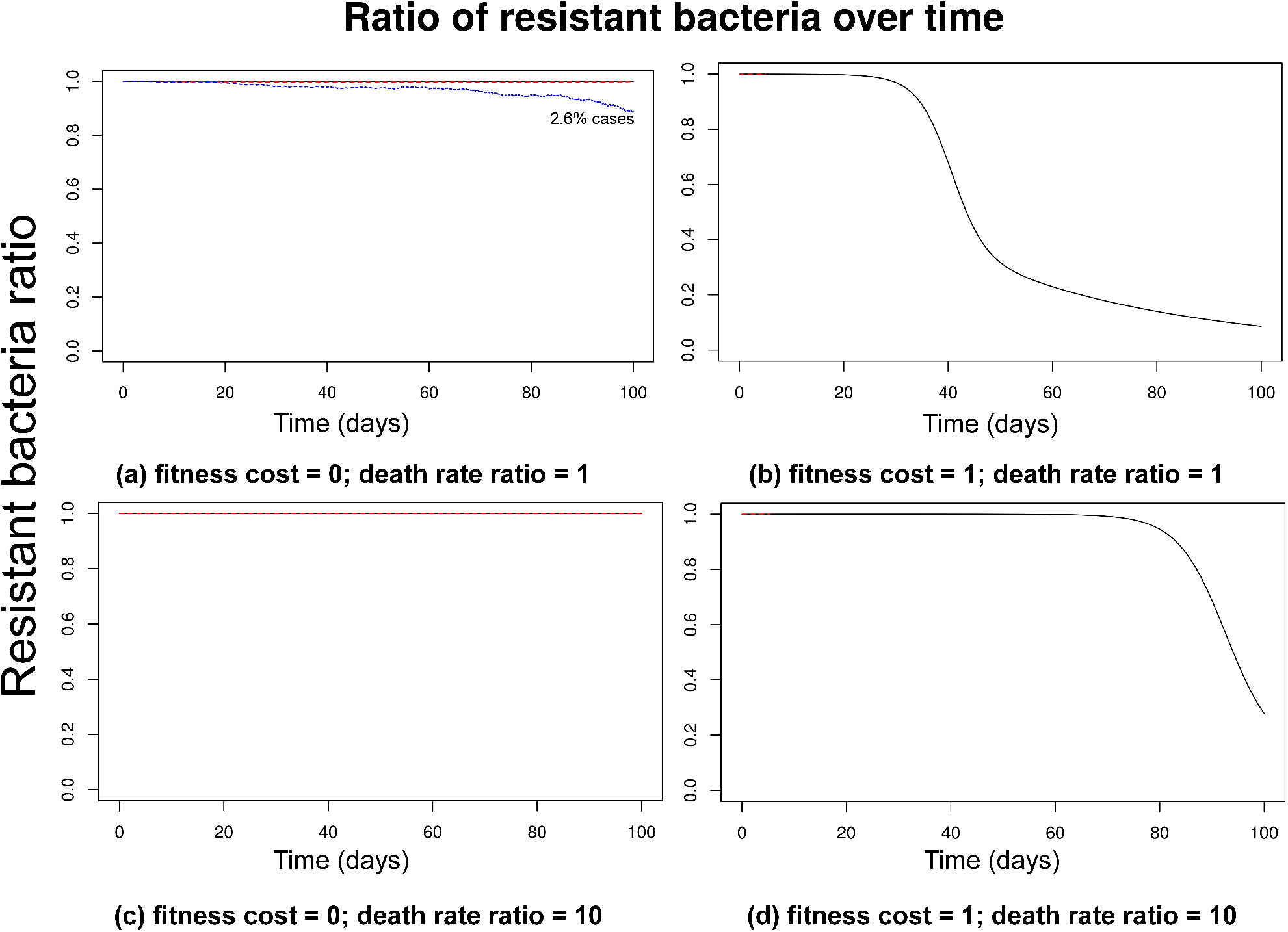
Time series of proportion of resistant bacteria for the bounding values of fitness cost and death rate ratio. Time series curves up to 100 days for select values of fitness cost and death rate ratio of sensitive to resistant bacteria, both for deterministic (black) and stochastic (red and blue dashed) versions of the model. The four figures correspond to the bounding values for the ranges fitness cost and death rate ratio used in later simulations. With no fitness cost ((a) and (c)) deterministic version results in persistence of resistance, but there might be some loss (2.6% cases) in the stochastic model in the absence of selection due to drift ((a)). With high fitness cost ((b) and (d)), there is loss of resistance, with the time for loss dependant on death rate ratio. The stochastic version in this scenario, however, leads to loss of both sensitive and resistant bacteria.

### Effect of toxicity and fitness cost on persistence of resistance

In order to evaluate persistence of resistance due to co-selection, we carried out a sensitivity analysis for two parameters, metal toxicity and fitness cost for plasmid carriage, first using the deterministic version of the model. We measured the time for the resistant population to drop from 99.32% to 50% (Fig 2). The dominant outcome for higher levels of metal toxicity (i.e. higher concentrations) and lower levels of fitness cost is persistence of resistance (grey zone in Fig 2). The coloured zone represents those simulations where resistance is lost, ranging from blue (rapid loss) to red (slower loss).

**Fig 2.**
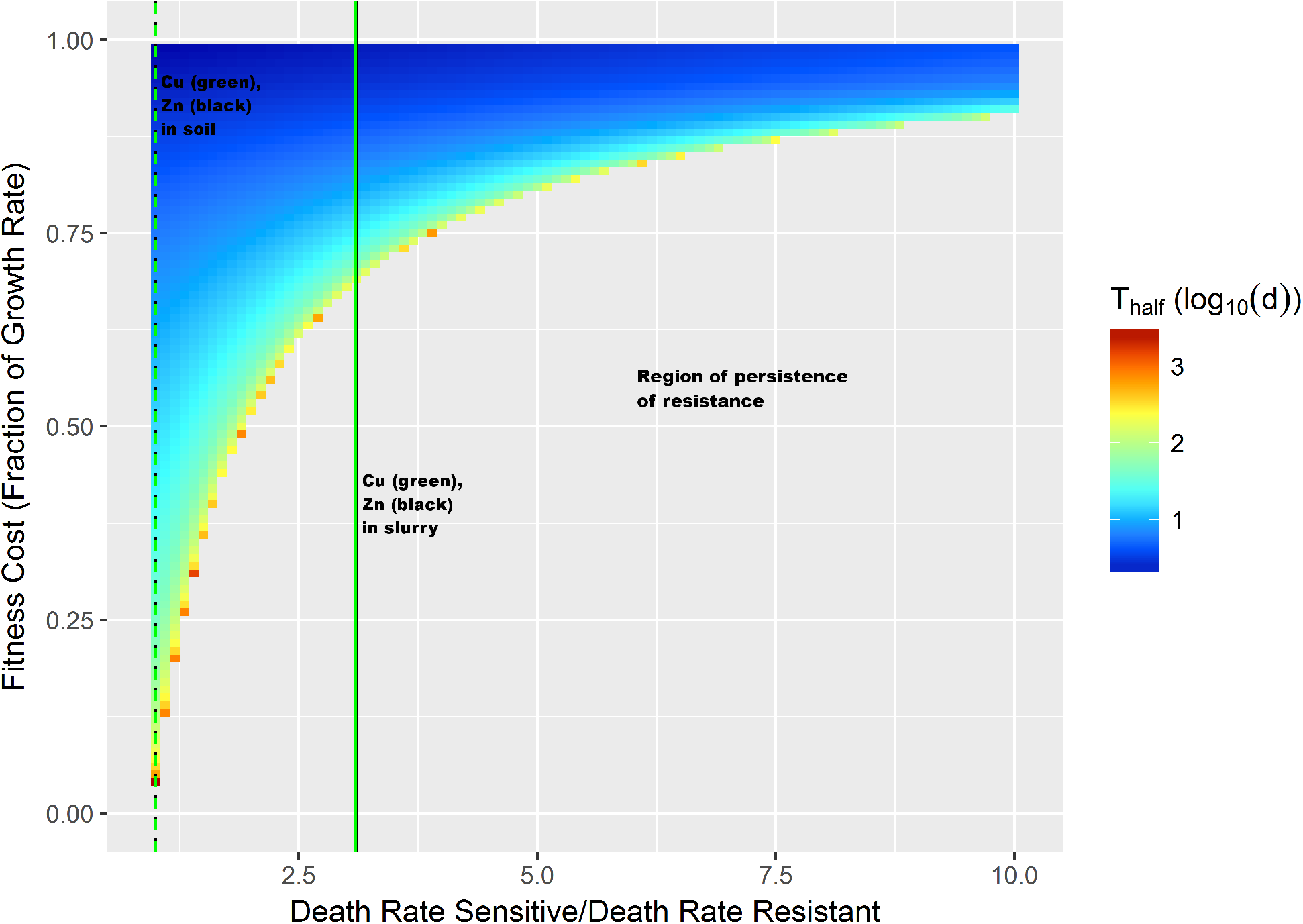
Sensitivity analysis of the deterministic model. The figure shows the number of days for the resistant population to drop to 50% total population from the same starting point of 99.32% resistant population, in the absence of antibiotic. The x-axis represents metal toxicity in terms of ratio of sensitive against resistant bacterial death rate. As can be seen, co-selection pressure causes persistence of resistance at low fitness cost and high metal toxicity. Loss of resistance is seen only with high fitness cost or with no metal-ratio of death rates equal to 1. The vertical lines represent the death rate ratio for copper (green) and zinc (black) concentrations in the example environments of slurry (solid lines) and slurry amended soil (dashed lines at almost identical x-coordinates) as measured by the method mentioned before. For both metals, the observed concentrations lead to similar death rate ratios, with higher chance of persistence in slurry than slurry amended soil.

A key feature to note is the steep rise in the boundary between the two zones: as the level of toxicity increases, the minimum fitness cost required for loss of resistance also increases sharply. This suggests that co-selection can occur even at low metal concentrations.

Compared to the deterministic model with a single outcome for a set of parameter values, a stochastic model may result in different outcomes on repeated runs with same parameter values. Thus, in order to assess the impact of stochasticity, the probability of different outcomes was coded as different colours on the RGB scale, with red denoting resistant bacteria at more than 50%, blue denoting resistant bacteria less than 50% and green denoting loss of all bacteria (Fig 3). Thus, a mix of these colours at a point signifies that the same parameter combination resulted in different outcomes. As can be seen in Fig 3(a), after 100 days there is a greater chance of persistence (red) rather than loss (blue), whereas after a 1000 days (i.e. 3 years) or 10^5^ days (chosen as a very long period to allow the model to equilibriate), there was a clear pattern of resistance loss or persistence, similar to the pattern for the deterministic model. Thus, the outcomes in Fig. 1 with loss of resistance in less than 100 days, have a low probability, as inferred from Fig 3(a). Even then, the fitness cost required for such loss, is higher than typical costs of stable plasmids [43, 47, 48].

**Fig 3.**
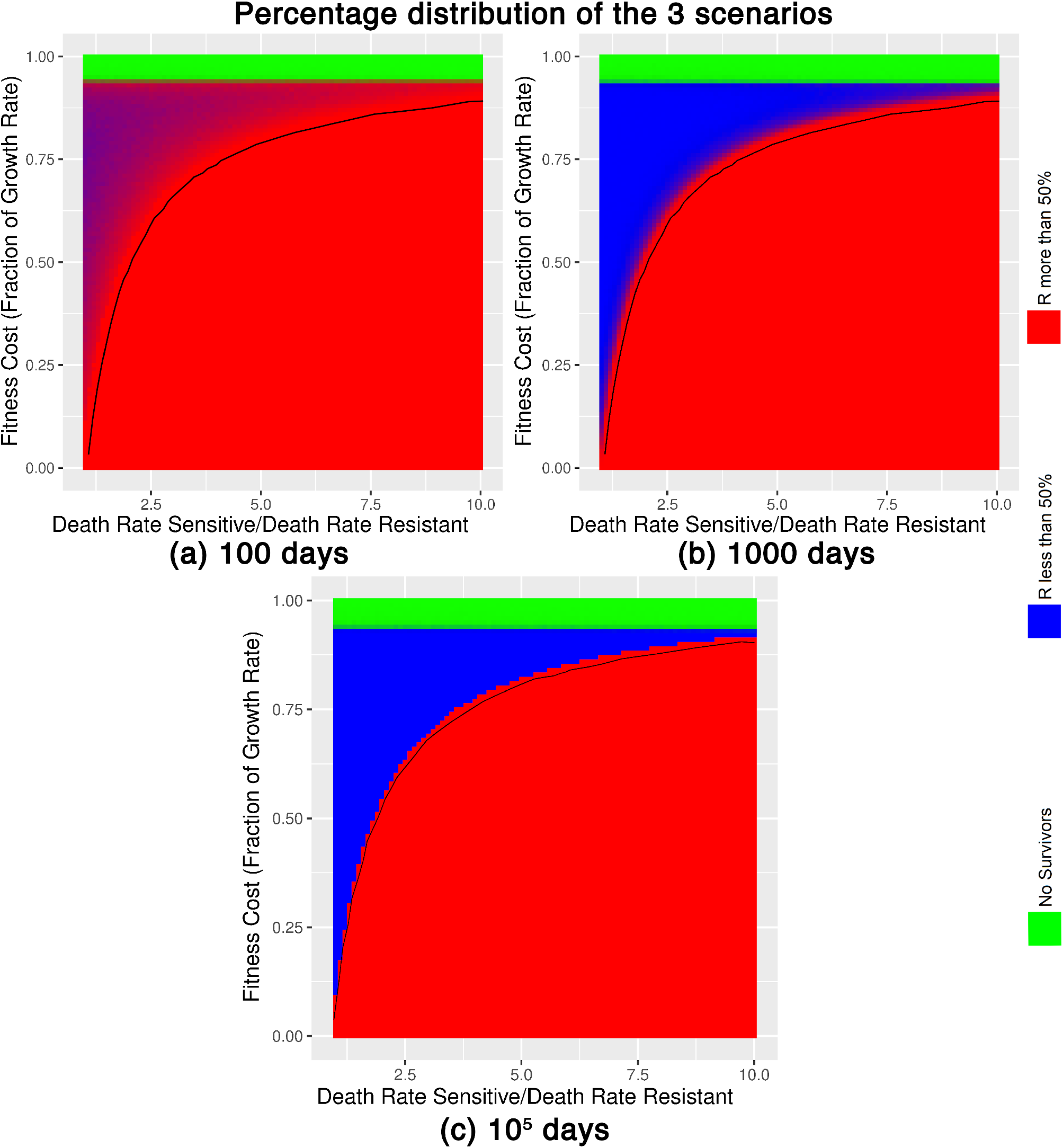
Sensitivity analysis of the stochastic model. Percentage distribution of three outcomes occurring in the stochastic simulations, with each outcome is represented by a different colour (red: persistence of resistance; blue: loss of resistance; green: total extinction). The three panels show different time points in the simulations, 100 days, 1000 days (i.e. 3 years) and 100,000 days (to reach steady state). In the short term, there is a greater probability of persistence of resistance as after 100 days, cases with relatively high fitness cost show very little loss of resistance but after 1000 days, we see a similar pattern as seen for the ODE simulations (black line). The black line corresponds to the edge of persistence region of Fig 2.

Our previous work highlighted the importance of gene transfer rate on spread of resistance [28]. Therefore, to assess the robustness of the outcomes shown in Fig 2 to changes in the conjugation rate parameter, we conducted similar sensitivity analyses with conjugation frequency 10 times higher (Fig. S1) and lower (Fig. S2). The result with lower conjugation frequency is almost identical, probably because conjugation is not an important factor when resistance is already high. On the other hand, with increased conjugation frequency, there is increased persistence of resistance. However, the default conjugation frequency (0.001 h^−1^) is already at the upper end of observed values, so higher values are unlikely to be realistic. Therefore, we are confident that the results shown are robust to variations in transfer rate. We also varied the value of probability of plasmid loss due to segregation, showing that a higher loss probability speeds up loss of resistance (Fig. S3), whereas a lower loss probability has similar outcome (Fig. S4). Again, higher rates of plasmid loss are not likely to be especially relevant because of plasmid addiction systems, and so we are confident that our results are robust to variations in this parameter too. Changing growth rate had no effect on the output (data not shown).

### Estimating minimum co-selective concentrations (MCSCs)

We presented model outcomes in terms of the ratio of death rates at different metal concentrations (Fig 2); these can be used to predict minimal co-selection concentrations for specific metal, bacterial and plasmid combinations, using a metal toxicity model (Eq. (3)), and knowledge of the fitness cost of carriage. For these calculations, we used a fitness cost of 0.25, which is within the reasonable range of expected values [43, 47, 48], although this could be reduced to produce more stringent MCSCs. Thus, the death rate ratio selected was 1.25. Using the estimated parameter values, we calculated the MCSC value based on this ratio. The data is presented in Table 3 for copper, zinc, mercury, lead and silver using *E. coli* as an exemplar.

**Table 3.** Metal MCSC and measured concentrations.

| Metal | MCSC (mg/L) | Farm Soil (mg/kg) | Dairy Slurry (mg/L) | ALS Environmental soil guidelines (mg/kg) |
| --- | --- | --- | --- | --- |
| Copper (Cu) | 5.5 | 0.068 | 22.32 | 80 (5.0<pH<5.4)<br>200 (pH≥7.1) |
| Zinc (Zn) | 1.6 | 0.16 | 32.16 | 200 (5.0<pH<5.4)<br>450 (pH≥7.1) |
| Mercury (Hg) | 0.0156 | NA | NA | 1 (pH>5.0) |
| Lead (Pb) | 21.5 | NA | 0.047 | 300 (pH>5.0) |
| Silver (Ag) | 0.152 | NA | 0.00026 | NA |

The MCSC threshold is calculated by assuming death rate ratio of 1.25, which corresponds to fitness cost of 0.25 for predicted loss of resistance in Fig. 2. A metal concentration above this threshold is predicted to provide co-selective pressure and lead to persistence of resistance. With our farm example data, concentrations of copper and zinc are predicted to be coselective in the slurry, but not slurry amended soil. Measured concentrations of lead and silver in the slurry are predicted not to be coselective.

In reference to our example environments, measured zinc concentrations are 32.16 mg/L (slurry) and 0.3 mg/L (soil), and copper concentrations are 22.32 mg/L (slurry) and 0.17 mg/L (soil). Therefore the metal concentration in slurry is above the MCSC for both metals, hence this will be classified as a co-selective environment; however, the metal concentration is below the MCSC for both metals in slurry-amended soil, so that would not be a co-selective environment. Similarly, the measured concentrations of lead and silver in slurry are both below the MCSC threshold and so these metals are not predicted to be co-selective.

## Discussion

Several studies have shown that there is a correlation between presence of metals and abundance of antibiotic resistance genes (ARGs) in soil, including in Scotland [51], Western Autralia [52] and urban soils from Belfast area [53]. These correlations are indicative of co-selection due to metal presence, but do not prove a causal link. The model described in this study provides a mechanistic insight into the different factors which drive co-selection. Although both deterministic and stochastic versions were defined with similar assumptions and parameter values, the inherent assumptions about the biological processes in each methodology are different and hence might lead to different results. The deterministic version shows that loss of resistance genes or resistant bacteria is only possible in low toxicity (lower death rate ratio) environment, or, in the rare case of cost prohibitive plasmids (high fitness cost). Most AMR phenomena are observed in large scale environments such as guts, tanks, fields, farms, hospitals [54], and the deterministic model can provide a reasonable approximation for prediction of the behaviour of large scale populations if they are, or can be considered to be, well-mixed. However, most environments are comprised of smaller, diverse, microscopic bacterial communities, and so deterministic models may be less realistic; thus stochasticity can play an important role [32]. The stochastic model in this work leads to a similar general conclusion of the effects of metal toxicity and fitness cost, but stochasticity leads to a greater chance of persistence over a shorter (reasonable) time-scale of antibiotic absence.

Importantly, the models we developed can be used to predict minimal co-selective concentrations (MCSCs) for transition metals. These predictions can help inform metal concentration thresholds for environments in which antibiotic resistant bacteria are likely to be present - provided other organisms don’t have toxicity values much lower than predicted MCSCs. Animal agriculture is a prime example due to both metal and antibiotic use. The MCSC determined here is low compared to the permissible concentrations set by established guidelines. For example, the ALS Environmental guideline for soil concentrations has the maximum permissible value for copper and zinc set at 80 mg/kg and 200 mg/kg, respectively, for pH between 5.0-5.4 and 200 mg/kg and 450 mg/kg respectively at pH of 7.1 or higher [55]. The same report gives the concentrations for mercury and lead at 1 mg/kg and 300 mg/kg, respectively, for pH 5.0 or higher. Similarly, Seiler and Berendonk [13] suggest an MCSC for soil of 117 mg/kg fresh weight for copper. Our study would indicate that current guidelines provided for soil metal concentrations are not sufficiently stringent and might provide a co-selective environment.

On the other hand, guidelines for maximum permissible concentration (MPC) for water - either freshwater, saltwater or groundwater (but not dirty water) - suggest extremely low metal concentrations. Taking the example of the report by National Institute of Public Health and the Environment Bilthoven, The Netherlands, we see that MPCs provided for copper are 1.5, 1.4 and 2.4 *μ*g/L [56], which is 1000 fold lower than the MCSC value estimated by the model. This might be due to more sensitive toxicity levels of other organisms found in these environments or the use of these sources for drinkable water. Similar values are reported for other metals as well. These figures are very similar to the MCMCs suggested by Seiler and Berendonk [13] for copper and zinc (1.5 *μ*g/L and 19.61 *μ*g/L respectively), but these concentrations may be difficult to apply to dirty water (e.g. slurry).

The current model does not explicitly account for changes in bioavailability of metal, which may cause change in toxicity values. For example, Cr^3+^ ions are less toxic than chromate and which form they occur in is dependant on environmental conditions [57]. Thus, this speciation (physiochemical form of metal) can affect the toxicity of the metals. Also, determination of element bio-availability remains unpredictable and contingent on adsorption dynamics, absorption within soil particles, flocculation, ion exchange, precipitation and complexation reactions. While classical geochemical Pourbaix relationships can provide insight about possible interactions based on pH and Eh (redox), a lot remains dependent on surface character and affinities, especially soil organic matter, water, salinity and temperature. Elevations in pH tend to reduce solubility, and the presence of carbon dioxide tends to promote carbonate precipitation; Eh reductions enhance sulfide precipitation; whereas salinity (or presence of multiple ions) tends to mobilize the metals.

The BCR483 extraction (NH_4_NO_3_) provides a good representation of trace element mobility [58], and represent the bio-availability from the sludge amended soils. The acid/peroxide extractions represent oxidizable forms and probably over-estimate metal lability, but does reflect the fraction associated with organic carbon, which can be highly dynamic in terms of complexation and solubility. Copper availability tends to be highly dependent on organic matter content, to the extent that kd values for Cu^2+^ tended to be independent of pH (when >5.5) and driven by organic carbon [59]. While sludge amendments can enhance organic matter content in soils, and Cu binding [60, 61], the presence of dissolved organic matter can mobilize the copper [62]. Lead strongly binds to organic matter, especially humic at pH >4. Shifts between anoxia and oxic conditions may induce periods of solubility, but remain reactive to sulfide precipitation. Solubility values tend to be associated with pH under low carbon and sulfur environmental matrices; in case here, it is likely to be associated with the nature of the organic matter. Zinc can become insoluble with sulfide at reducing conditions and can form relatively insoluble carbonate precipitates at higher pH. However, lability of zinc best correlates with total zinc concentrations [63], rather than precipitation reactions. Dissociation reactions are similar whether applied as sludge or as a salt [60]. Silver in the environment, while extremely toxic, is relatively insoluble. It strongly binds to organic matter and oxides within the soils, to the point that desorption is considered negligible.

The toxicity model defined in this study, uses the bioavailable metal concentration values and the Hill equation to calculate the death rate. While this empirical approach fits the data, more mechanistic approaches that take into account the details discussed above could be appropriate. Moreover, the toxicity values that are reported by Ivask *et al.* [33] are not based on the environment and do not take into account the interaction between different metals or metal and other biocides.

Despite these assumptions, the model can be applied to a large number of environments, with relatively minor changes. However, in more complex environments, spatial heterogeneity and stochasticity may become more important [32]. Another complexity that is missing and might provide further insight into the role of metals towards co-selection is the inclusion of multiple metals. Environmental contaminants occurring in a mixture is an observed and quantified norm. Thus, the presence of multiple metals might affect their bioavailability/toxicity. Data exists to show the correlation between contaminant mixture and ARGs [64], but, this only proves that there is co-occurrence of multiple toxicants and a higher abundance of ARGs. A mechanistic understanding of co-selection due to multiple toxicants could provide interesting further results. Similar data, if available for other co-selective agents like biocides, QACS, etc. can be used to calculate the death rate and thus understand their potential for antibiotic resistance co-selection.

In conclusion, our model shows that co-selective pressure can maintain antibiotic resistance in microbial communities, even in the absence of antibiotic. It provides a general approach for setting appropriate standards for transition metal contamination, especially for environments where antibiotic resistance is likely to be important, e.g. in livestock farming, and monitoring those environments against those standards. It also implicates the importance of developing technologies for removing metals from such environments [65].

## Supporting information

**Fig. S1.**
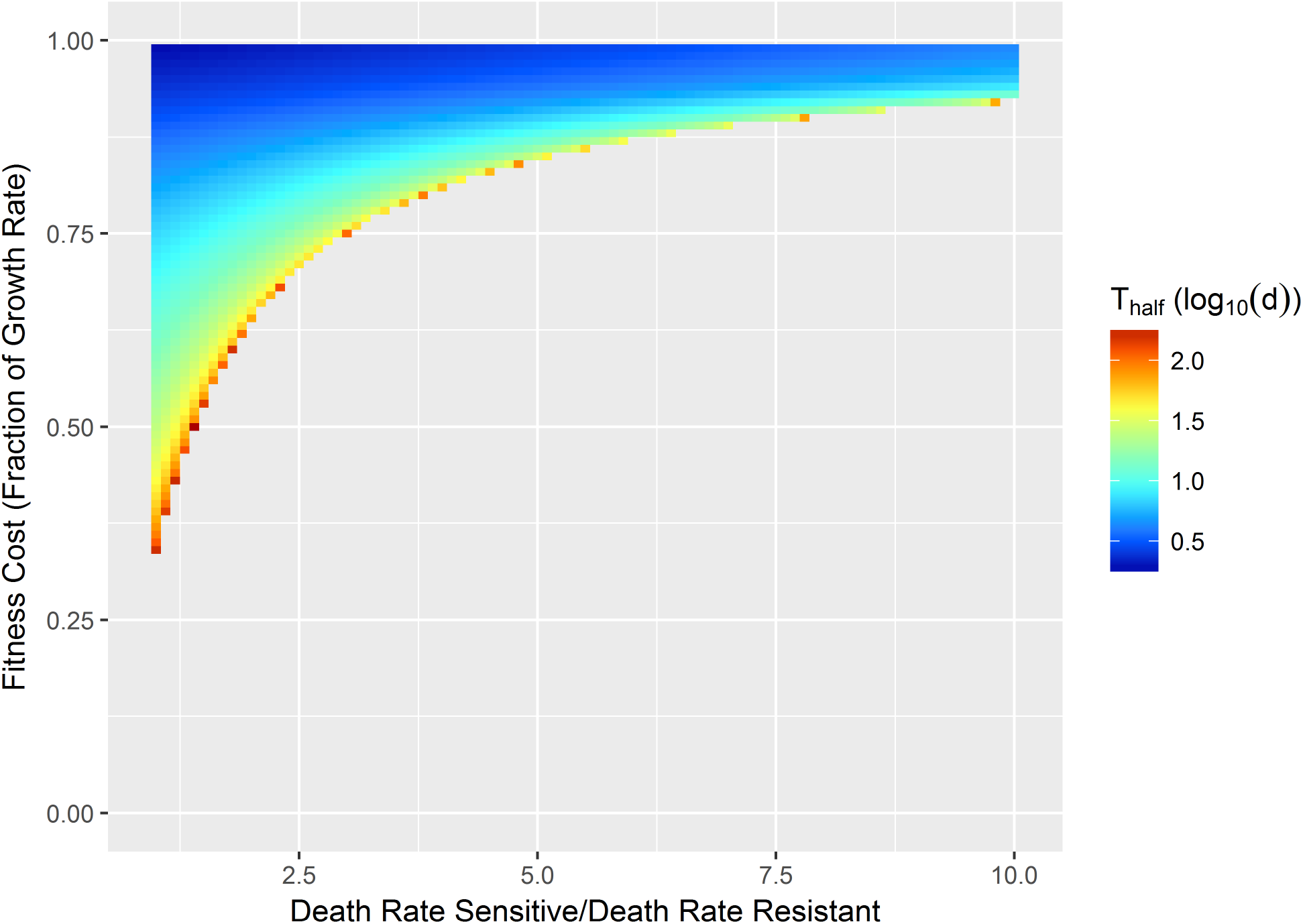
Conjugation rate is 0.01, i.e., 10 times more than for Fig 2. We can see this leads to greater chances of persistence of resistance, even in situations with no selection pressure.

**Fig. S2.**
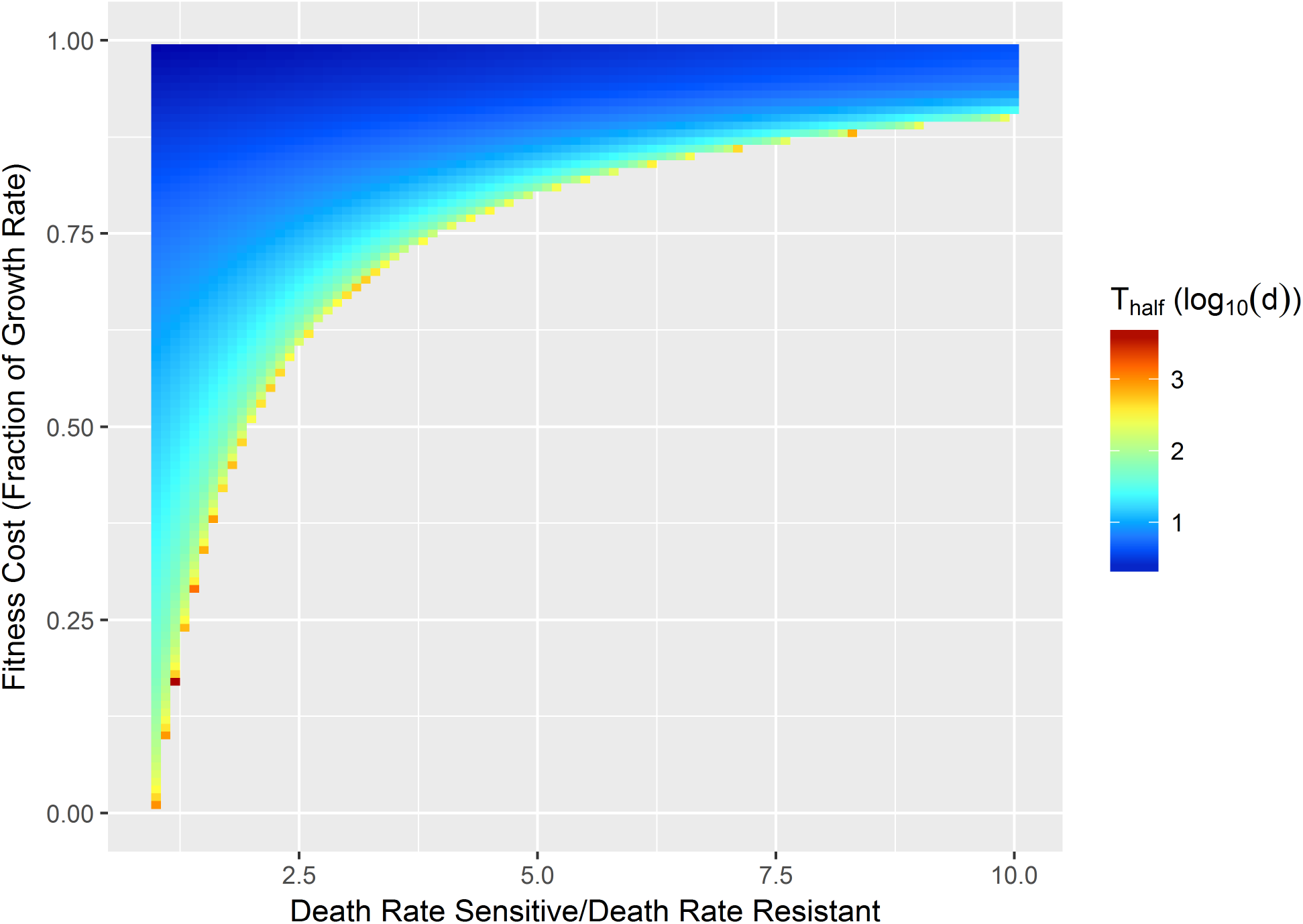
Conjugation rate of 0.0001 (10 times lower). Shows very little difference compared to Fig 2.

**Fig. S3.**
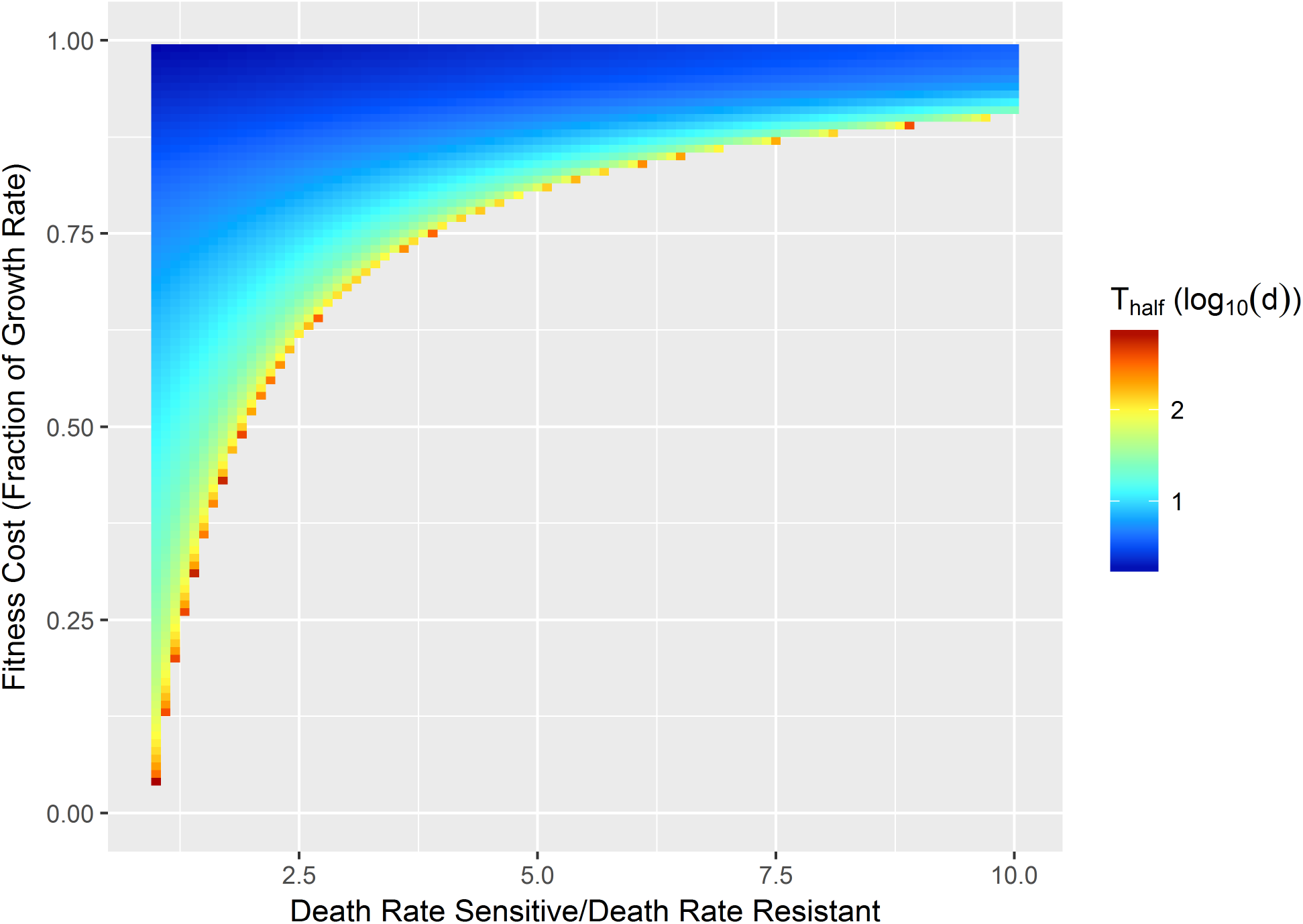
10 times higher probability of loss of resistance carrying plasmid due to segregation (0.00144). The affect is seen in the time required for loss of resistance. Most of the situation that lead to persistence are not affected, but where loss is likely the time required for loss of 50% of resistance from population is reduced by approximately a factor of 10, as the red colour corresponds to 10^3^ days, when before it was 10^4^ days.

**Fig. S4.**
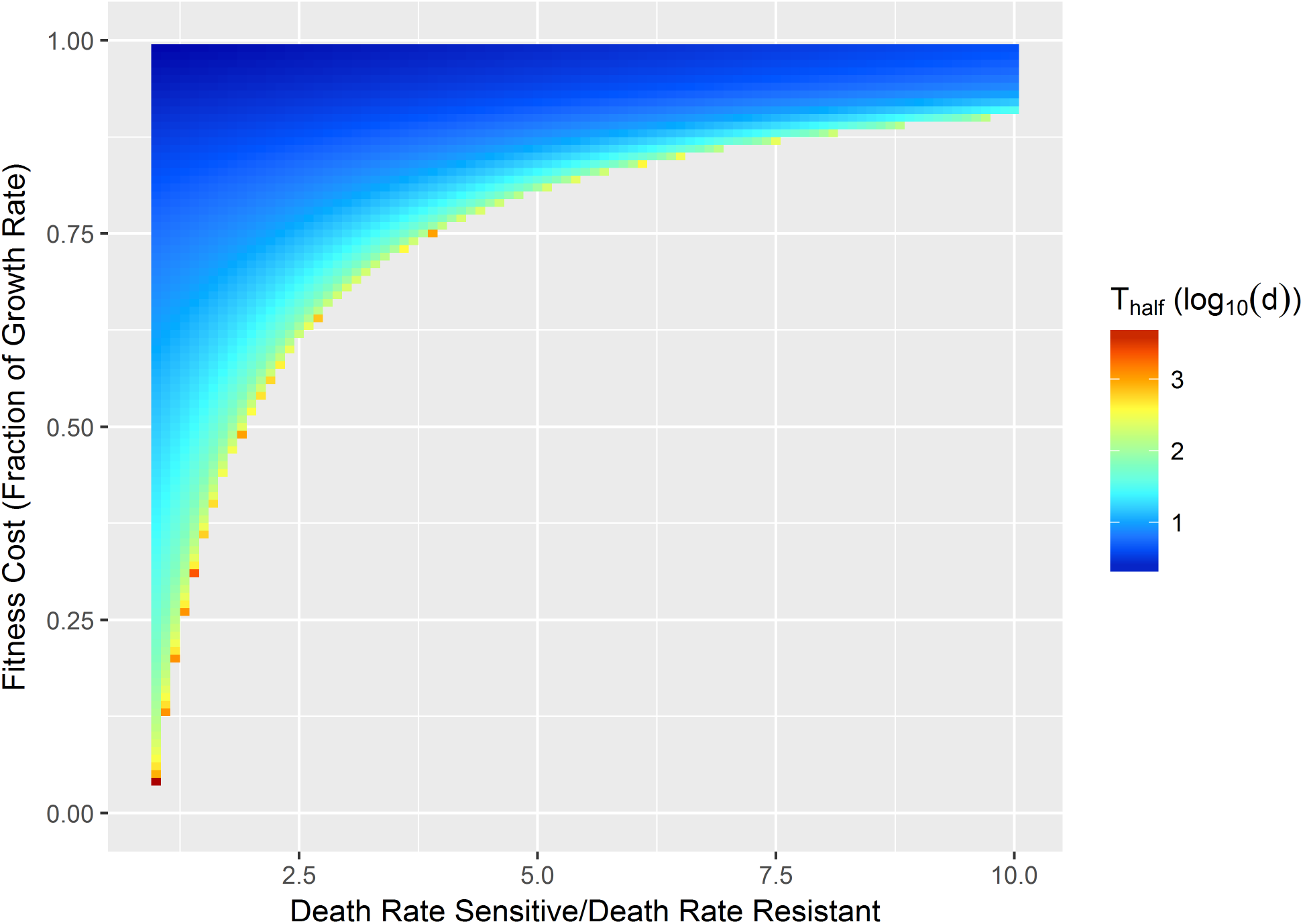
10 times lower probability of loss of plasmid due to segregation (0.0000144). Similar results to that seen in Fig 2

**Fig. S5.**
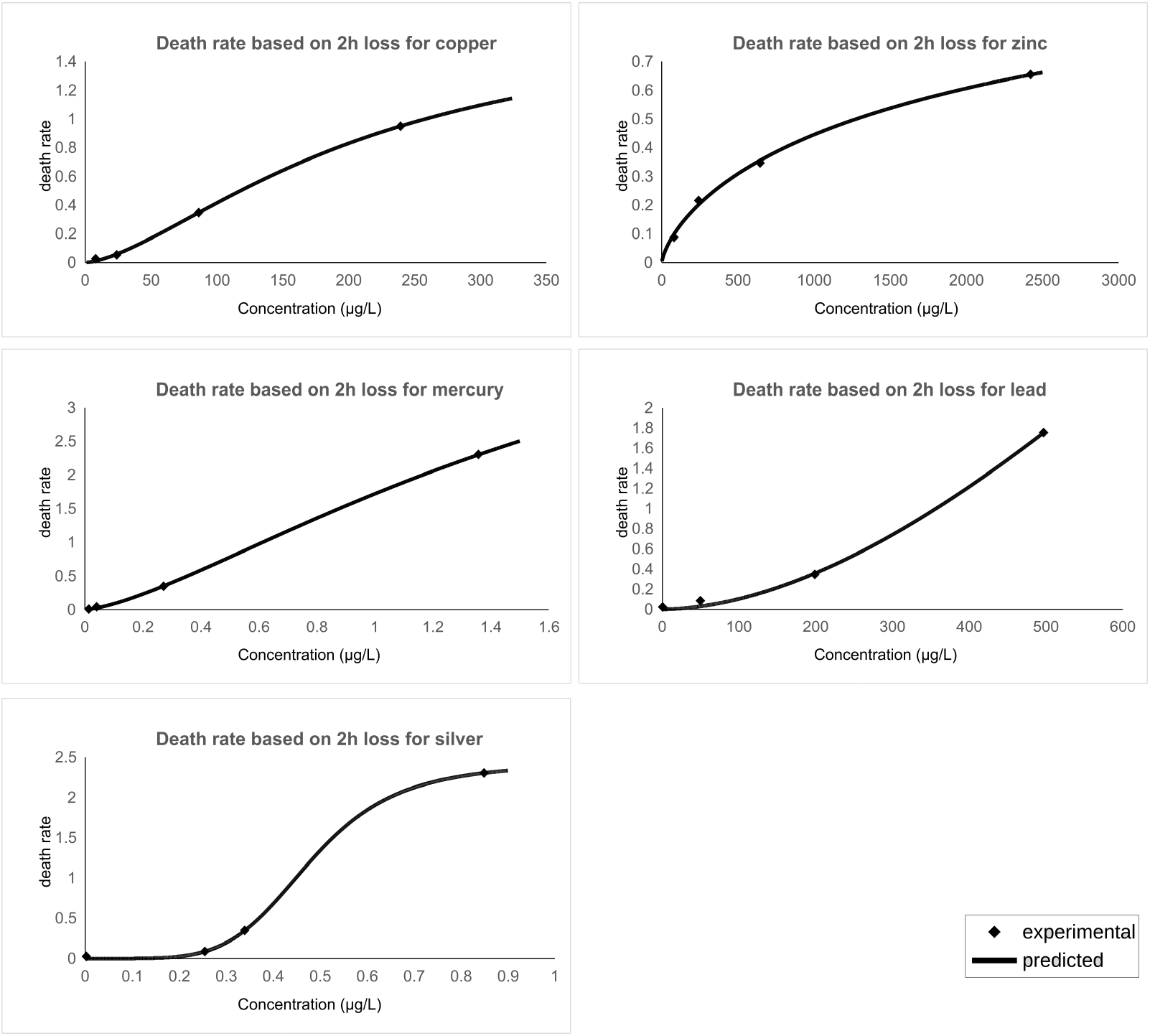
Fits of metal toxicity data. Figure shows the predicted death rate (line) and experimental death rate values (points) for each of the metals considered.

## Acknowledgments

Sankalp Arya was funded by a University of Nottingham Vice Chancellor’s Scholarship. Alexander Williams was funded by that Natural and Environmental Sciences Soil Training and Research Studentship. We thank Scott Young for facilitating the ICP-MS work.

## Notes

### Competing Interest Statement

The authors have declared no competing interest.

